# Polymorphisms Predicting Phylogeny in Hepatitis B Virus (HBV)

**DOI:** 10.1101/2022.07.05.498824

**Authors:** José Lourenço, Anna L McNaughton, Caitlin Pley, Uri Obolski, Sunetra Gupta, Philippa C Matthews

**Author notes:** Shared first co-authorship (José Lourenço, Anna L McNaughton).

## Abstract

Hepatitis B viruses (HBV) are compact viruses with circular genomes of ∼3.2kb in length. Four genes (HBx, Core, Surface and Polymerase) generating seven products are encoded on overlapping reading frames. Ten HBV genotypes have been characterised (A-J), which may account for differences in transmission, outcomes of infection, and treatment response. However, HBV genotyping is rarely undertaken, and sequencing remains inaccessible in many settings. We used a machine learning approach based on random forest algorithms (RFA) to assess which amino acid (aa) sites in the genome are most informative for determining genotype. We downloaded 5496 genome-length HBV sequences from a public database, excluding recombinant sequences, regions with conserved indels, and genotypes I/J. Each gene was separately translated into aa, and the proteins concatenated into a single sequence (length 1614aa). Using RFA, we searched for aa sites predictive of genotype, and assessed co-variation among the sites with a Mutual Information (MI)-based method. We were able to discriminate confidently between genotypes A-H using 10 aa sites. 5/10 sites were identified in Polymerase (Pol), of which 4/5 were in the spacer domain, and a single site in reverse transcriptase. A further 4/10 sites were located in Surface protein, and a single site in HBx. There were no informative sites in Core. Properties of the aa were generally not conserved between genotypes at informative sites. Co-variation analysis identified 55 pairs of highly-linked sites. Three RFA-identified sites were represented across all pairs (two sites in spacer, and one in HBx). Residues that co-vary with these sites are concentrated in the small HBV surface gene. We also observe a cluster of sites adjacent to the Surface promoter region that co-vary with a spacer residue. Overall, we have shown that RFA analysis is a powerful tool for identifying aa sites that predict HBV lineage, with an unexpectedly high number of such sites in the spacer domain, which has conventionally been viewed as unimportant for structure or function. Our results improve ease of genotype prediction from limited regions of HBV sequence, and may have implications for understanding HBV evolution and the role of the spacer domain.

## INTRODUCTION

Hepatitis B virus (HBV) is the prototype virus of the *hepadnaviridae* family, a family of small, circular viruses with partially double-stranded (ds)DNA genomes of ∼3.2kb in length(1). The viral genome encodes seven proteins within four genes – HBx, Core, Polymerase and Surface (Table 1; Figure S1) – together with associated regulatory elements(2), arranged in a series of overlapping reading frames. This genome structure imposes constraints on selection, acting as a stabilising selective force during replication(3, 4), and accounting for a reduced nucleotide substitution rate in overlapping regions (approximately 40% lower than in the non-overlapping regions)(5).

**Table 1.** Summary of HBV genes and proteins, and their roles and functions

| Gene | Protein(s) | Roles and function |
| --- | --- | --- |
| X | X | • Small regulatory protein (154aa)<br>• Role in subversion of host restriction factors<br>• Transactivating properties that are implicated in oncogenesis(33). |
| Core (C) | Pre-core (e-antigen); Core | • Post-translational processing to derive capsid protein and e-antigen<br>• Roles in intracellular trafficking and stabilisation of covalently closed circular (ccc)DNA<br>• Soluble e-antigen is secreted into blood, can cross the placenta; acts as an immune tolerogen |
| Polymerase (Pol) | Polymerase | • Four distinct domains: terminal protein (TP); spacer; reverse transcriptase (RT); RNase H.<br>• Takes up approximately two-thirds of the genome(34).<br>• RT and RNase H domains show homology to HIV proteins, and some HIV nucleos(t)ide inhibitors can be used for HBV treatment(34).<br>• TP and spacer domains are unique, and no known homologues have been identified to date(34). |
| Surface (S) | Short (S)<br>Medium (Pre-S2 + S)<br>Long (Pre-S1 + Pre-S2 + S)<br>surface proteins | • External envelope<br>• Receptor binding domains<br>• Surface epitopes neutralised by vaccine-mediated or naturally arising antibodies<br>• Produced in excess, with a potential role as a tolerogen/immunological decoy.<br>• Gene is completely overlapped by polymerase, representing the longest known gene overlap of any animal virus(35). |

HBV DNA genomes are copied via RNA intermediates by means of an error-prone viral reverse transcriptase (RT) enzyme(6), driving an evolutionary rate that is higher than would be expected for a DNA virus with a high density of overlapping reading frames(1). The resulting genetic diversity is the basis for the classification of HBV into ten genotypes (gt), defined by ≥7.5% nucleotide divergence(7), and designated gt-A-I, along with an unusual recombinant putative gt-J (showing similarity to gt-C and gibbon *orthohepadnavirus*)(8). Genotypes are further classified into subgenotypes based on ≥4% divergence(7). There is variation in the number of subgenotypes per genotype, ranging from >10 subgenotypes in gt-C(9) (reflecting its status as the oldest lineage(5)), to just a single subtype in gt-E, -G and -H.

To date, HBV sequencing (and genotyping) is not recommended at baseline by clinical guidelines and is not routinely undertaken to inform patient care, as there has been insufficient evidence to support its role in informing surveillance or determining treatment courses(10). However, as the pool of HBV sequence data expands, alongside linked clinical metadata, progressive insights are emerging into associations between sequence heterogeneity (including genotype, insertions, deletions and polymorphisms) and different clinical phenotypes including treatment response and disease outcomes(10, 11).

Machine learning approaches are frequently applied to omics-based data, including transcriptomics and proteomics(12). We set out to apply a machine learning approach based on a random forest algorithm informed by full-length HBV sequences. Our aim was to identify genome regions through which genotype can be predicted and to cast light on the selection pressures that determine HBV genetic population structure.

## METHODS

Random forest algorithms (RFA) are a type of decision tree-based analysis, providing a relatively hypothesis-free approach to interrogating complex data sets. The method has been applied widely, including in host tropism studies in influenza(13), to identify molecules inhibiting flaviviruses(14), to analyse mutational fitness effects in picornaviruses(15), and in the identification of genes related to immunogenicity and pathogenicity in *Streptococcus pneumoniae* infection(16, 17).

Briefly, this study included nucleotide alignments (n=5496) of HBV genotypes A-H(9) (Table S1). Recombinant sequences were excluded from the analysis, as were genotypes I and J which are recombinant in origin. Each of the overlapping HBV genes was separately translated into amino acid (aa) sequences, which were then concatenated into a single sequence for each genome (total length 1614 aa, Figure S1B). Residues were numbered and reported using X02763 (gt-A) as a reference sequence, as is convention in the field(9). The RFA pipeline, as detailed in the supplementary methods and Figures S3 and S4, was then applied to the concatenated HBV sequences, using the known genotype of each sequence as the classification variable and aa sites as predictive variables, in search of a parsimonious number of sites that maximised prediction of sample genotype (feature selection).

To address the impact of site co-variation on feature selection, we quantified amino acid co-variation among all pairs of sites in the HBV genome using a Mutual Information (MI) approach as previously applied to *Plasmodium falciparum* sequence data(18). A full description of the methods can be found in the supplementary material.

## RESULTS

### HBV genotypes can be distinguished through 10 amino acid sites

The machine learning approach discriminated confidently between HBV gt-A-H using just 10 amino acid sites (Figure 1). Half of these sites (5/10) were identified in Pol, with four in the spacer region of Pol, and a single site in the reverse transcriptase (RT) domain. A further 4/10 sites were located in the Surface protein, particularly in pre-S1 (2/4), and a single site was identified in HBx. The majority of the sites (9/10) were in overlapping regions (the single site in RT being the exception), with the pre-S1/spacer overlap accounting for 6/10 sites. None of the 10 sites identified were in Core protein.

**Figure 1:**
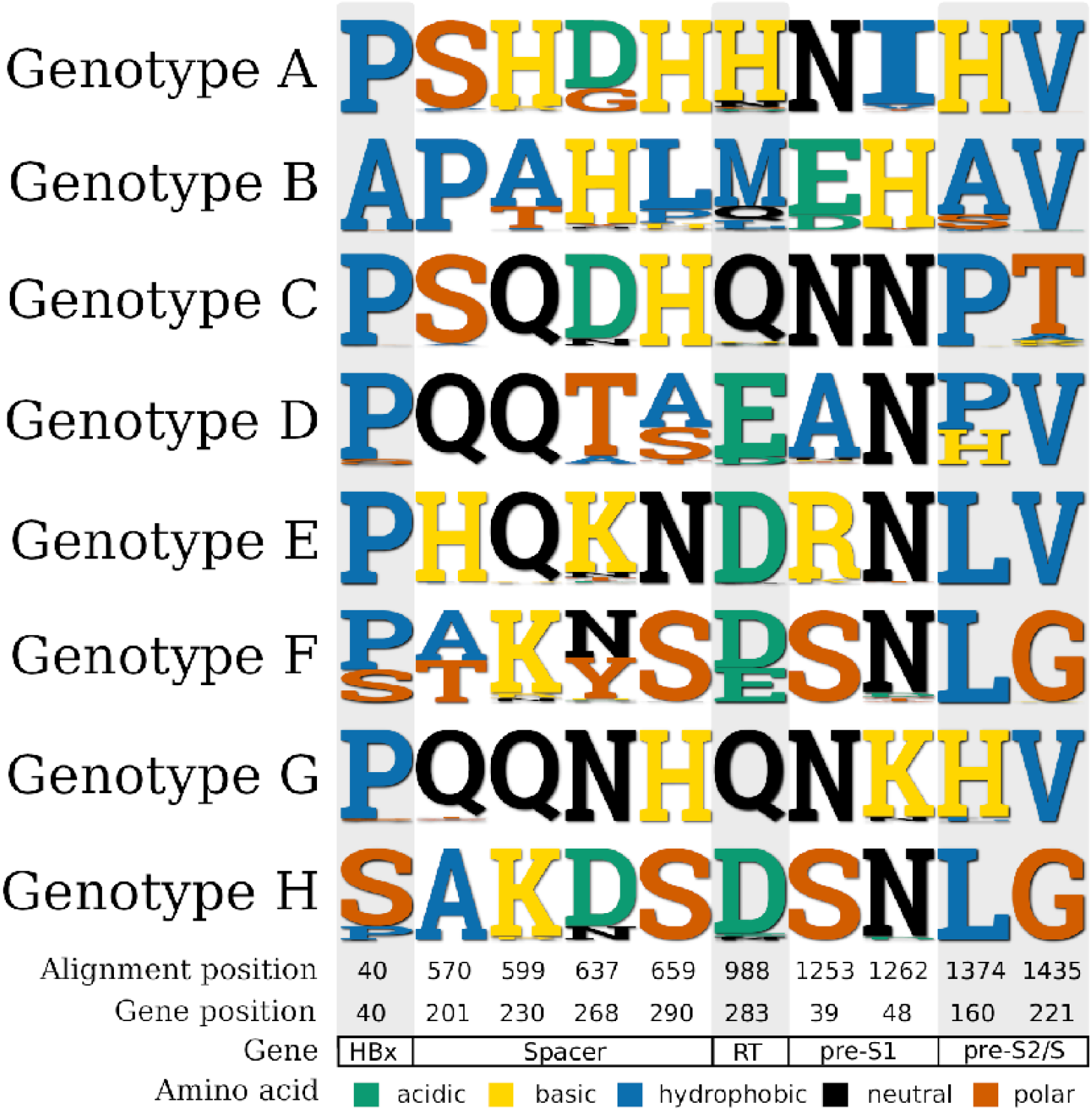
Top 10 amino acid sites discriminating between HBV genotypes, and the residues found at the sites in each genotype. Residues have been coloured according to their properties (key at bottom of the figure). Position of the sites is given in the concatenated amino construct used for analysis (Figure S1) and the equivalent locations in each gene are given at the foot of the figure. In Pol, residues in spacer are given assuming the first amino acid is at the start of the terminal protein, and sites within the reverse transcriptase (RT) domain are counted separately, as is convention in the field (Table S3).

We classified the amino acid sites based on chemical properties (Figure 1). Properties were generally not conserved across the genotypes at informative sites, with the exception of HBx-40 which was almost always hydrophobic apart from in gt-F/H. This observation suggests that there may be consistent selection pressure to maintain different chemical properties between genotypes, and that the sites are not located in key regions required for host interactions, as this would typically require functional conservation.

### General location of the top genotype-informative sites

We considered the top 50 most informative sites to determine whether this changed the distribution throughout the genome compared to the top 10 sites. The distribution of these sites within the genome remained comparable. In particular, amino acid 40 in HBx remained the only informative site in HBx, 28/50 (56%) were located in Polymerase, with 16/28 of these sites located in the spacer domain. The Surface protein contained 21/50 sites. The majority of sites were identified in overlapping regions of the genome, with a low number of sites in the TP, RT and RNAse H domains of the Polymerase polyprotein (Table S2, Figure S2).

### Sites defining genotypes

Although 10 sites were sufficient to discriminate confidently between all genotypes considered, the majority of sites identified were conserved within each genotype, albeit with a few sites presenting variation at the subgenotype level. For example, gt-B sequences could be identified by a single site, 40A in HBx, with all other genotypes having 40P/S (Figures 1, S5). The 988H residue (283H in RT) was also key for identifying gt-A (Figures 1, 2). Other sites were polymorphic within a particular genotype, but genotype-specificity could be distinguished by the *absence* of particular residues (e.g. non-V/G at site 1435 (221 in surface) indicates gt-C) (Figures 1, S6). The close evolutionary history of gt-F and H could be seen by homology at many of the top-10 sites (Figure 1), with site 637 (87 in spacer domain) demonstrating the clearest discrimination between gt-F (637N/Y) and gt-H (637D) (Figure S5). Sites 40 and 570 also showed differences in the distribution of amino acids between gt-F and gt-H.

**Figure 2:**
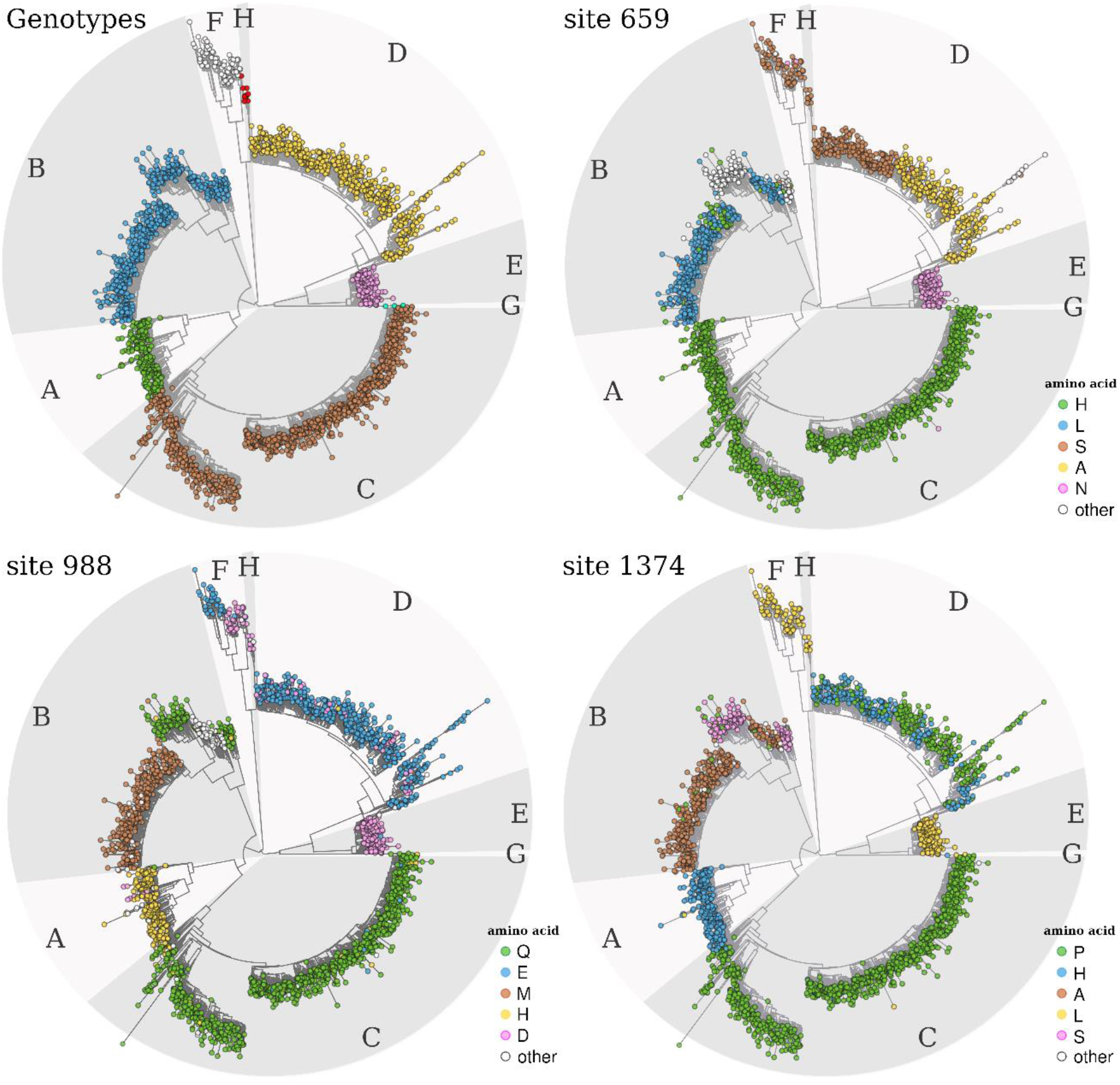
Phylogenetic trees showing overall genotype lineage, and distribution of three exemplar amino acid sites that predict lineage. Maximum likelihood phylogenetic trees were available to download as a part of the online resource from which we obtained nucleotide sequences(9). The top left tree highlights the different HBV genotypes (A to H) with capital letters and with shaded, alternating, lighter and darker grey areas. For the trees of sites 659, 988 and 1374, nodes are coloured on the basis of the amino acid residue at each site (inner legends) using an inhouse R script based on the R package ‘Analyses of Phylogenetics and Evolution’ (ape v5.4 (36)). On each tree, only the top five most frequent amino acids are presented, with the rest under the category “other”. Phylogenies for the other 7 top-10 sites are shown in Supplementary Figures S5 and S6.

### Sites defining subgenotypes

A number of the top-10 informative sites were also highly discriminatory for some HBV subgenotypes, including p40-P/S (gt-F), p599-A/T (gt-B), p637-D/G (gt-A) and p659-A/S (gt-D) (Figures 2, S5 and S6). Differences in the amino acids selected at some of the sites in gt-D, in particular p40-S and p1253-R (Figures S5-6), also support the designation of gt-D5 as a unique subtype(9). These sequences cluster distantly from other gt-D sequences on a long branch and show strong geographical clustering, with all sequences isolated from India and Bangladesh.

### Co-varying sites with top 10 informative sites

When using Random Forests, high co-variation between two predictor variables can result in their importance for classification being shared and thus penalized (relative to other predictor variables). Within our pipeline, this could have resulted in the exclusion of pairs of sites that present high co-variation, or the exclusion of single sites that had high co-variation with the top 10 selected sites. To address these possibilities, we first used Mutual Information theory to quantify co-variation between all pairs of amino acid sites across the genome (see Supplementary Text for full details), and found that the vast majority of site pairs present low co-variation (Figure S7).

Among the 55 site pairs with the highest co-variation (Table S4), all included at least one site from the top 10 most informative sites that discriminate genotype, thus ruling out the possibility that pairs of sites that discriminate genotype and that present high co-variation were excluded in our Random Forest approach. Of the top 10 sites, only three featured in 55 site pairs with the highest co-variation (HBx site 40, and spacer sites p599 and p637) presenting varying degrees of co-variation with a range of sites across the genome (Table S4, Figure S8). These other sites, although not in the top 10 list of sites that discriminate genotype, could still be of biological interest. For example, we find that HBx 40 is highly co-variable with aa227 in Pol (p596) and a series of amino acids at the start of small-HBs (p1403, p1404, p1406, p1411; corresponding to aa15-23), which also co-varied with site p599 but to a lesser degree. The highest co-variations were found between p599 and both HBx aa39 (p39) and pre-S1 aa97 (p1311), which were also found to intermediately co-vary with HBx 40. Reasons for the strong association between this site in HBx, spacer and the start of small-HBs are unclear. The majority of past work on HBx interactions has focused on interactions with host proteins rather than considering influences on other viral proteins(19, 20).

In comparison, site p637 (aa268 in Pol) had the lowest degrees of co-variation with other sites, but nonetheless presented a varied list of connections, showing associations with two sites in core (aa100 and 123; p254 and 277 respectively) and a cluster of closely-located sites in Pol (p632, p633, p636). This cluster of sites in Pol overlaps a regulatory region in the nucleotide code adjacent to a ‘CCAAT’ box, known to be the S-promoter region(21).

## DISCUSSION

### HBV genotypes can be defined by 10 key sites

Our analysis demonstrates that HBV genetic population structure can be determined from as few as 10 amino acid sites across the viral proteome (Figure 1, Table S3). Four of the top-10 sites were identified by previous studies as informative for HBV genotyping (Table S3). Analysing the aa sequences individually by protein has enabled us to determine which residues are key, avoiding difficulties in interpretation that could otherwise arise as a result of the overlapping genome structure. Our analysis further suggests that Core is uninformative for distinguishing HBV genotypes (Table S2). This is in keeping with a high conservation rate of >75% of amino acid sites(22), as expected for a highly structural capsid protein which also plays diverse roles in the viral replication cycle. HBx was also found to be a relatively uninformative region of the HBV genome, with a single site identified in the top-50 most informative sites (Table S2).

### Informative sites are concentrated in the spacer domain

The spacer domain, which spans aa 184-348, is an intrinsically disordered protein and poorly conserved region of Pol, unique to hepadnaviridae. Previous literature has shown that the spacer domain can tolerate significant deletions and insertions without a significant impact on polymerase function(23, 24).

The unexpected clustering of sites that predict genotype in the spacer domain indicates that whilst the domain retains a considerable amount of plasticity, this is highly lineage-specific rather than stochastic. Other studies have also found that spacer mutations are relevant in distinguishing between simian hepatitis B viruses(25), as well as human HBV genotypes(9, 26, 27) and subgenotypes(28–30). Importantly the four top-10 sites we identified in spacer map to regions previously identified as useful for lineage distinction(24). This suggests selection pressures may be acting to conserve genotype-specific sequence within spacer. Furthermore, it substantiates the hypothesis that spacer plays a central role in the co-evolution of the overlapping P and S genes, potentially related to selection pressure from antiviral drugs, vaccines and the host immune response(24). In addition to encoding regions within proteins, the promoter region for the RNA transcript encoding medium- and small-HBs is present in the pre-S1/spacer overlap region. Mutation of the spacer region is therefore likely to interfere with the generation of M-/S-HBs transcripts, the biological significance of which is unclear(31). Current models are poorly equipped to study this, suggesting our understanding of the role of spacer may be limited by the tools used to analyse its function.

### Limitations of the methodology

Recombinant sequences, combining two or more different HBV genotypes, were intentionally excluded from this analysis. As such, the short list of 10 sites that can discriminate genotypes is not expected to be able to adequately classify recombinant samples(32). Since our algorithm used the amino acid sequences to compare isolates, synonymous mutations are not considered in the analysis. As several regions of the HBV genome contain promoter regions, such as the well-described basal core promoter of pre-core, synonymous changes in the DNA sequence would have important functional effects. Conservation of the nucleotide sequence in these regions would therefore also be key and may be lineage-specific.

## CONCLUSIONS

We present the observation that HBV can be reliably genotyped using information from as few as ten sites, and for the distinction of some genotypes by a single site. This is of potential practical importance if a genotype identification is desired but limited sequence data are available. Our finding that discriminatory sites are concentrated in spacer underlines the role and evolutionary importance of the spacer domain in the viral polymerase. With emerging importance of genotypes in HBV disease outcomes, quick approaches to genotyping from short fragments of sequence data may be of increasing practical utility, particularly in low-resource settings. Furthermore, describing the impact of selection pressure at different sites in the genome can provide insights into viral evolution, and potentially contribute to mechanistic insights regarding viral persistence and pathogenesis.

## Acknowledgements and Funding

ALM was supported by a National Institute for Health Research (NIHR) Research Capability Funding grant [award CO-CIN-01]. JL was supported by a Principal Investigator research contract by FCiências.ID (Associação para a Investigação e Desenvolvimento de Ciências) of the Faculty of Sciences at the University of Lisbon. PCM is funded by the Wellcome Trust (grant ref. 110110/Z/15/Z), University College London Hospitals NIHR Biomedical Research Centre (BRC), and The Francis Crick Institute.

