## Supplementary data file for "Polymorphisms Predicting Phylogeny in Hepatitis B Virus (HBV)"

#### Supplementary Methods

##### Data sources

We downloaded genome-length alignments of HBV sequences genotyped A-H from HBVdb(1) in June 2018, obtaining a total of 5496 sequences. Recombinant sequences were excluded from the analysis, along with two additional isolates (FJ023674 and FR714503) identified as gt-C recombinants. Genotypes I and J were also excluded as genotype I is recombinant in origin (comprising sequences from gt-A, -C and -G)(2) and only a single isolate of genotype J has ever been isolated(3). The number of sequences in each genotype is listed in Table S1. The phylogenetic trees included in this analysis were adapted from an open-access online resource on HBV reference sequences(4).

##### Numbering of HBV genome for analysis

We followed convention in the field, using the gt-A sequence X02763 as a numbering reference and defining nucleotide (nt)1 in the middle of an EcoR1 restriction site in the P/S overlapping region(4)(Figure S1A). In order to analyse the amino acid sites in each HBV gene independently, without confounding by overlapping regions, we translated each of the genes from nucleotide into amino acid separately. The translated amino acid sequences were then concatenated into a single sequence, comprising the X, core, polymerase and surface genes (total 1614 amino acids based on a gt-A reference, Figure S1B). Regions with conserved deletions or insertions were excluded from the analysis in all sequences. All deletions removed resulted in proteins that were still in-frame and the locations of them are described by McNaughton *et al*, 2020(4).

##### Random forest classification approach

Using random forests, we developed a pipeline that searches for the top M most informative amino acid sites, in terms of classifying genotypes. M can be any number of amino acid sites and the pipeline stops when that number is reached or when it can no longer guarantee to reach M. This pipeline is described in detail below, and a visualisation is included as Figure S3.

(1) Start by running a series of RFAs with the intention of finding a set of amino acid sites that are generally highly informative and much smaller in number than the length of our sequence alignment.

(1.a) Create K training / test folds. Folds are data partitions, created by randomly sampling ~80% of the dataset for training and the remainder ~20% for testing. Sampling is done such that the number of samples of each genotype is the same between sets.

(1.b) For each pair of training / test K folds run an RFA and memorize the top N most informative sites.

(1.c) Next, create a set of N* most informative sites as the intersection of the N top sites of each K RFA run. The set N* is of minimum size N, but can be larger since each K RFA run may result in some of the N top sites being different from the other K RFA runs.

(2) Perform feature (site) selection among the N* set, by attempting to iteratively remove the less informative sites while minimizing prediction error. First, set a minimum number of sites of interest, M (the number for which we wish to stop). Then, run the RFA with the N* set of sites and order them by their resulting importance.

(2.a) Remove the site with lowest importance, resulting in a proposed new set N*.

(2.b) Run the RFA with the new N* set and check for an error heuristic (see below for definition).

(2.b.i) If the heuristic is not violated, then the site is permanently removed from the N* set. If the size of the set N* is still larger than M, the process jumps to step (a); otherwise the process stops.

(2.b.ii) If the heuristic is violated, the last removed site is returned to the N* set and the site with the next lowest importance is removed. The process jumps to (b).

If the removal of all sites in N* is attempted consecutively without any removal, the process stops.

Within the pipeline, we took the simple heuristic of minimizing the miss rate (false negative) across genotypes. This is done by strictly not allowing the miss rate (false negative) of any genotype to be above 2.5%. Thus, in step 2.b of the pipeline, if the miss rate (false negative) of all genotypes is below 2.5%, step 2.b.i is performed, otherwise step 2.bii. In the context of the 8 classes (genotypes), the miss rate (false negative) of each is calculated as follows:

predicted as genotype

| known genotype | A | B | C | D | E | F | G | H |
|---|---|---|---|---|---|---|---|---|
| A | pAA | pAB | pAC | pAD | pAE | pAF | pAG | pAH |
| B | pBA | pBB | pBC | pBD | pBE | pBF | pBG | pBH |
| C | pCA | pCB | pCC | pCD | pCE | pCF | pCG | pCH |
| D | pDA | pDB | pDC | pDD | pDE | pDF | pDG | pDH |
| E | pEA | pEB | pEC | pED | pEE | pEF | pEG | pEH |
| F | pFA | pFB | pFC | pFD | pFE | pFF | pFG | pFH |
| G | pGA | pGB | pGC | pGD | pGE | pGF | pGG | pGH |
| H | pHA | pHB | pHC | pHD | pHE | pHF | pHG | pHH |

Miss rate (false negative) of genotype i = $Miss_i$ = $\frac{\sum_j \quad pij,i\neq j}{pii + \sum_j \quad pij,i\neq j}$, where pij is the proportion of samples of genotype i that were classified as j.

We use two measures of (classification) performance to present results: accuracy and multi-class area under the receiver operating characteristic curve (multi-class AUC).

Accuracy (ACC) is the proportion of samples correctly classified: $ACC_i = \frac{pii}{\sum_i \quad \sum_j \quad pij}$, where pij is the proportion of samples of genotype i that were classified as j. Multi-class AUC was calculated using multiple ROC curves as proposed by Hand and Till [38]. Visualisation of pipeline performance is provided in Figure S4.

#### Sites that co-vary with the top 10 sites selected by the Random Forest pipeline

When using RFA, an amino acid site's importance is a score of how important the site is to classify the genotype of a sample. If two sites are highly predictive of genotype but are also highly correlated with each other, they will present a "shared importance" score, reflecting that they hold redundant information. In the RFA pipeline we develop, it is theoretically possible that: (a) sites that are informative to classify genotype but that are highly correlated may end up not being selected in the top N sites, or (b) sites that have not been selected in the top N list may be highly correlated with selected sites and they could thus be sites of biological interest.

To assess these two possibilities, we used normalized Mutual Information (MI) to quantify site co-variation between every pair of sites in the sequence alignment. MI is an aggregate measure of entropy between two variables, varying between 0 and 1, with 1 representing full co-variation (every time a site changes amino acid, so does the other site). We have previously applied MI to genetic sequences(5). The resulting MI distribution is long tailed (Figure S7A), with the vast majority of site pairs presenting low co-variation and a small number of site pairs presenting high co-variation. We considered a threshold of MI>0.6 to explore whether site pairs with high co-variation included sites in the top 10 list selected by the RFA pipeline.

This MI threshold resulted in 55 pairs of sites (Table S4). Among these, all contained at least one site from the top 10 list. This reassures us that when selecting the top 10 sites with the RFA pipeline, we have not penalized pairs of sites that have high co-variation to the point of disregarding such pairs (thus addressing point (a) above). Of the top 10 sites, only sites 40, 599 and 637 presented high co-variation levels with other sites in the alignment (Figure S7C-D), suggesting that these other sites hold redundant information that may be of biological interest. We describe these other sites in the main text and Table S3, thus addressing point (b) above.

#### Supplementary Tables

**Table S1: Number of full genome length HBV sequences in each genotype downloaded from HBVdb(4) and included in the analysis**

| HBV genotype | Number of sequences included in analysis | Number of sequences included in analysis |
|---|---|---|
| A | 755 | 794 |
| B | 1430 | 1450 |
| C | 1805 | 1830 |
| D | 856 | 882 |
| E | 246 | 250 |
| F | 225 | 226 |
| G | 36 | 38 |
| H | 26 | 26 |

**Table S2: List of 50 most informative sites.**

| Number of informative sites | HBx | Core | Pol | Surface |
|---|---|---|---|---|
| N=10 | 40 | n/a | 570 599 637 659 988 | 1253 1262 1374 1435 |
| N=20 | 40 | n/a | 570 599 624 625 637 659 666 839 988 | 1253 1262 1305 1322 1346 1374 1392 1435 1452 1547 |
| N=30 | 40 | n/a | 490 570 599 607 621 624 625 637 655 659 666 688 839 868 988 1092 | 1253 1262 1268 1305 1322 1346 1374 1392 1435 1452 1473 1531 1547 |
| N=40 | 40 | n/a | 452 490 570 599 607 610 621 624 625 637 639 655 659 660 666 688 771 839 868 988 1092 | 1253 1262 1268 1288 1305 1322 1329 1346 1374 1392 1433 1435 1444 1452 1473 1522 1531 1547 |
| N=50 | 40 | n/a | 439 445 452 490 570 585 599 602 607 610 621 624 625 637 639 655 658 659 660 666 683 688 771 838 839 868 988 1092 | 1253 1262 1265 1268 1288 1305 1322 1329 1346 1374 1392 1433 1435 1444 1452 1473 1501 1502 1522 1531 1547 |

**Table S3: Top-10 informative amino acid sites for determining HBV genotypes A-H identified using the RFA analysis.** Location of the sites in the concatenated construct used for analysis, the gene and domain/open reading frame (ORF) are provided.

| Position in concatenated construct | Gene | Protein aa site | Domain/ ORF | Domain/ ORF aa site | Further information |
|---|---|---|---|---|---|
| **40** | X | 40 | X | 40 | Located within ‘flexible loop region’ (residues 27–82); HBx site 40 has been previously identified as a site that is informative for genotyping(6) |
| **570** | P | 201 | Spacer | 20 | Nothing found |

| Position in concatenated construct | Gene | Protein aa site | Domain/ ORF | Domain/ ORF aa site | Further information |
|---|---|---|---|---|---|
| **599** | P | 230 | Spacer | 49 | Nothing found |
| **637** | P | 268 | Spacer | 87 | Nothing found |
| **659** | P | 290 | Spacer | 109 | Nothing found |
| **988** | P | 629 | RT | 283 | Nothing found |
| **1253** | S | 39 | S1 | 39 | Located within the NTCP-binding region of preS1 (aa2-48)(7); Neutralising antibodies can target region(8); Identified in previous genotyping studies based on pre-S1(9) |
| **1262** | S | 48 | S1 | 48 | Located within the the NTCP-binding region of preS1(aa2-48)(7); Identified in previous genotyping studies based on pre-S1(9) |
| **1374** | S | 160 | S2 | 41 | Identified in previous study as a key site in determining HBV serotype(10); Work with site-directed mutagenesis confirmed that Lys160 confers serotype *w* reactivity(11) |
| **1435** | S | 221 | S | 47 | Nothing found |

**Table S4: Top 55 site pairs with highest co-variation (MI>0.6) along the genome (all pairs of sites have one top 10 site that discriminates genotype).** Pairs of sites with very high co-variation are highlighted in orange by MI>0.8.

| Top 10 site discriminating genotype | Site presenting high co-variation | Location (gene/domain) | Mutual Information (degree of co-variation) |
|---|---|---|---|
| 40 | 583 | Pol/ spacer | 0.830347772449188 |
| 40 | 1328 | S/ pre-S1 | 0.829831434345946 |
| 40 | 1065 | Pol/ RT | 0.828633704636651 |
| 40 | 1432 | S/ surface | 0.827743343467526 |
| 40 | 1403 | S/ surface | 0.826907900110766 |
| 40 | 1411 | S/ surface | 0.826473553490255 |
| 40 | 596 | Pol/ spacer | 0.824939382569735 |
| 40 | 117 | HBx | 0.76673766529506 |
| 40 | 1404 | S/ surface | 0.759764763790752 |
| 40 | 817 | Pol/ RT | 0.704531874041169 |
| 40 | 1311 | S/ pre-S1 | 0.690291708728172 |
| 40 | 1406 | S/ surface | 0.678688383980139 |
| 40 | 39 | HBx | 0.66245335125875 |
| 40 | 706 | Pol/ spacer | 0.639127366833386 |
| 40 | 1355 | S/ pre-S2 | 0.637231787881651 |
| 40 | 569 | Pol/ spacer | 0.636954790343082 |
| 40 | 539 | Pol/ TP | 0.600260973512188 |
| 599 | 1311 | S/ pre-S1 | 0.88836302338456 |
| 599 | 39 | HBx | 0.885661154285354 |
| 599 | 706 | Pol/ spacer | 0.79119901897533 |
| 599 | 1355 | S/ pre-S2 | 0.788849670933397 |
| 599 | 583 | Pol/ spacer | 0.687558718839529 |
| 599 | 1432 | S/ surface | 0.683518612827625 |
| 599 | 1411 | S/ surface | 0.682376363037618 |
| 599 | 1065 | Pol/ RNAse | 0.680577777769191 |
| 599 | 1328 | S/ pre-S1 | 0.680200562467301 |
| 599 | 1403 | S/ surface | 0.674342417538883 |
| 599 | 596 | Pol/ spacer | 0.66775619591403 |
| 599 | 1371 | S/ pre-S2 | 0.654404578132584 |
| 599 | 117 | HBx | 0.620985334765634 |
| 599 | 1404 | S/ surface | 0.617844973891406 |
| 599 | 817 | Pol/ RT | 0.616344504959495 |
| 637 | 542 | Pol/ TP | 0.740581799342971 |
| 637 | 1192 | Pol/ TP | 0.739888327109659 |
| 637 | 1191 | Pol/ TP | 0.739571944411711 |
| 637 | 744 | Pol/ RT | 0.738483043208214 |
| 637 | 543 | Pol/ TP | 0.738179636789123 |

| Top 10 site discriminating genotype | Site presenting high co-variation | Location (gene/domain) | Mutual Information (degree of co-variation) |
|---|---|---|---|
| 637 | 562 | Pol/ spacer | 0.733980562647683 |
| 637 | 610 | Pol/ spacer | 0.730934749546506 |
| 637 | 1259 | S/ pre-S1 | 0.726699830812029 |
| 637 | 1212 | Pol/ TP | 0.71668888772881 |
| 637 | 277 | C | 0.673791585556711 |
| 637 | 1211 | Pol/ TP | 0.666992256184093 |
| 637 | 662 | Pol/ spacer | 0.661716806367433 |
| 637 | 434 | Pol/ TP | 0.661301456784865 |
| 637 | 254 | C | 0.656233802027875 |
| 637 | 432 | Pol/ TP | 0.642522541715073 |
| 637 | 615 | Pol/ spacer | 0.639253476138466 |
| 637 | 1264 | S/ pre-S2 | 0.638862208406785 |
| 637 | 636 | Pol/ spacer | 0.636239248939585 |
| 637 | 633 | Pol/ spacer | 0.635083322518339 |
| 637 | 1341 | S/ pre-S1 | 0.629140790565824 |
| 637 | 571 | Pol/ spacer | 0.626455257362722 |
| 637 | 632 | Pol/ spacer | 0.61610899673668 |
| 637 | 466 | Pol/ TP | 0.608188369453428 |

### Supplementary Figures

**Figure S1: HBV sequence map for analysis.** (A) Layout of HBV genome showing overlapping gene structure. The HBV genome encodes four open reading frames; HBx (X), Core (pre-C, C), Polymerase (P) and Surface (pre-S1, pre-S2, S). (B) Concatenated protein sequence generated from the amino acid translations of the HBV genes used for analysis. The gene and domain boundaries are given for Core, Polymerase and Surface, using the gt-A sequence X02763 as a numbering reference.

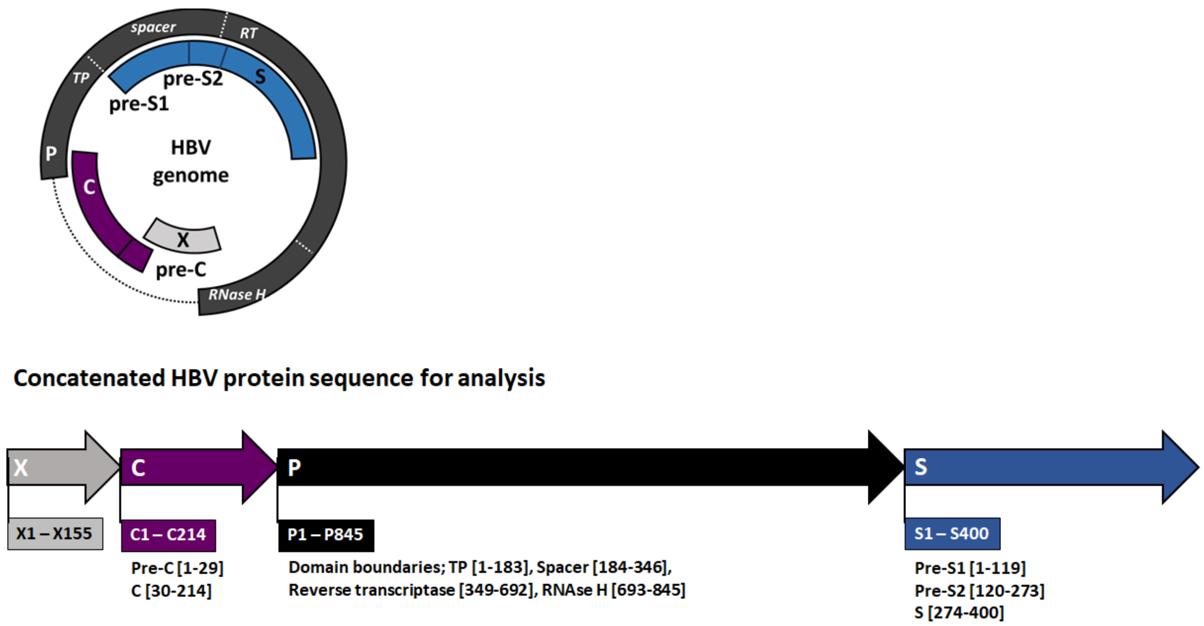


**Figure S2: Locations of the top-50 sites within the HBV genome (listed in Table S2).**

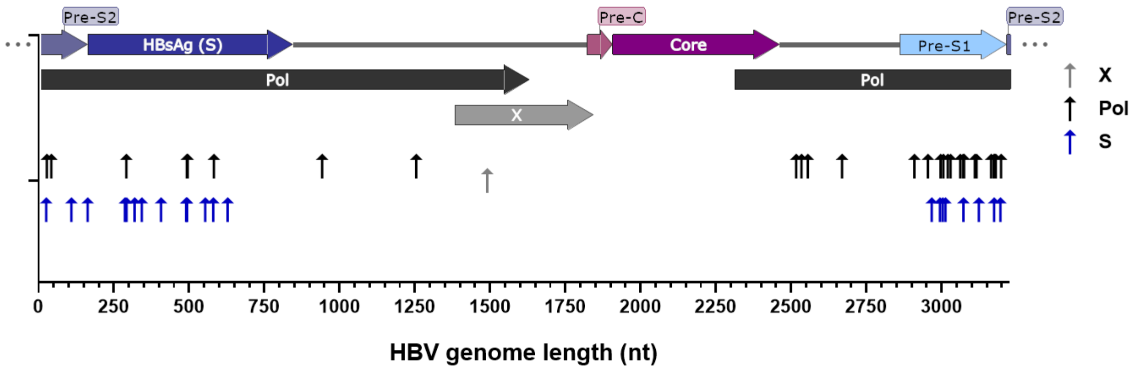

**Figure S3: Diagram of the pipeline.** Each shape represents a stage or action in the pipeline, and each arrow the flow between stages and actions. The sets of steps described in the main text (1.a-c; 2.a-2b.ii) are identified in the diagram.

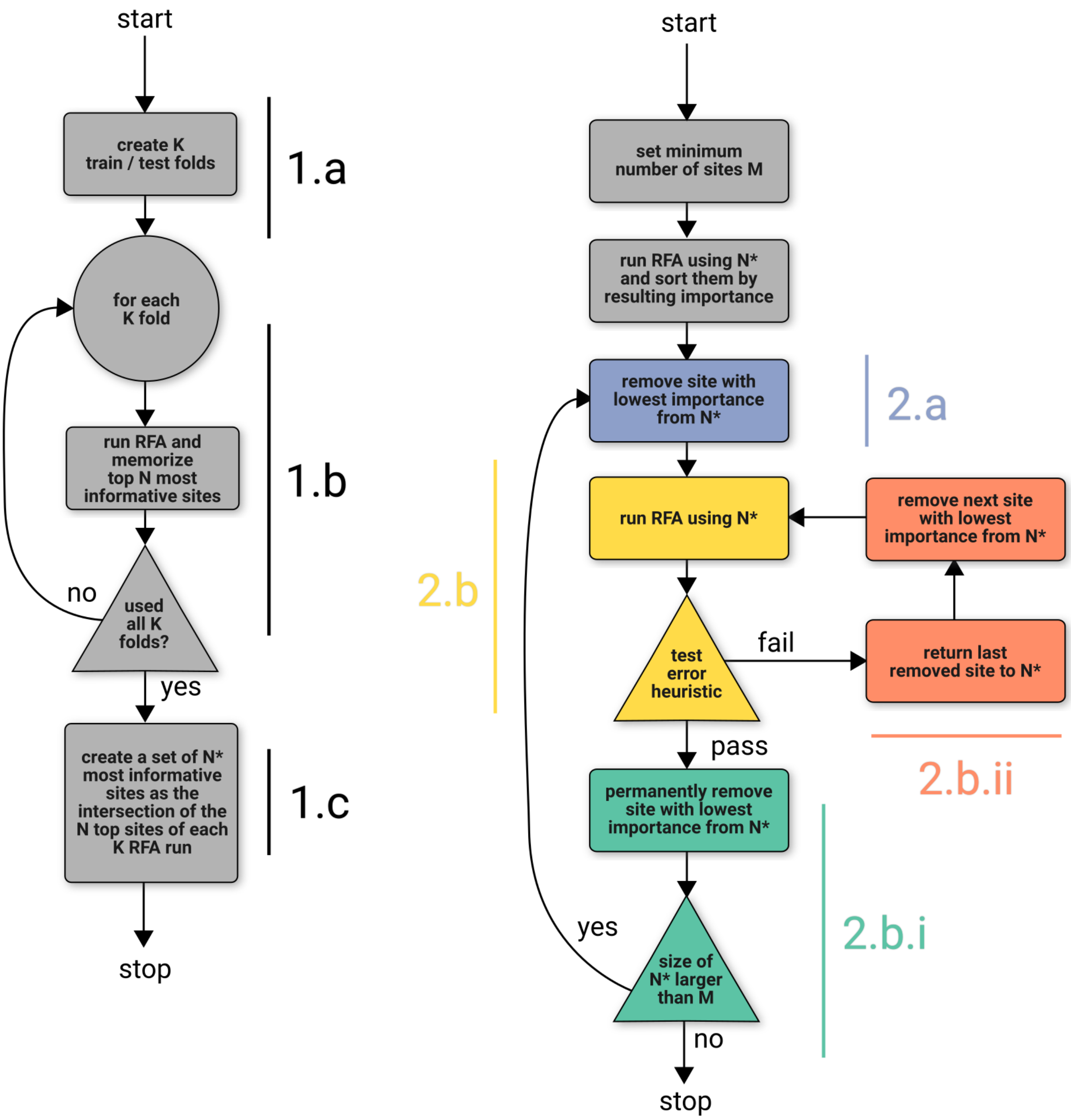

**Figure S4: Performance measures of the pipeline during feature selection.** Top row: Area under the curve, accuracy and miss rate given a number of sites (features) used for genotype classification (from left to right, respectively). Bottom row: Same as in top row but restricted to results starting at 40 sites. After N=10 the miss rate (false negative) of genotype F goes above the error heuristic, which is the stop condition for the pipeline. The resulting N=10 sites are the ones described in Tables S2,3.

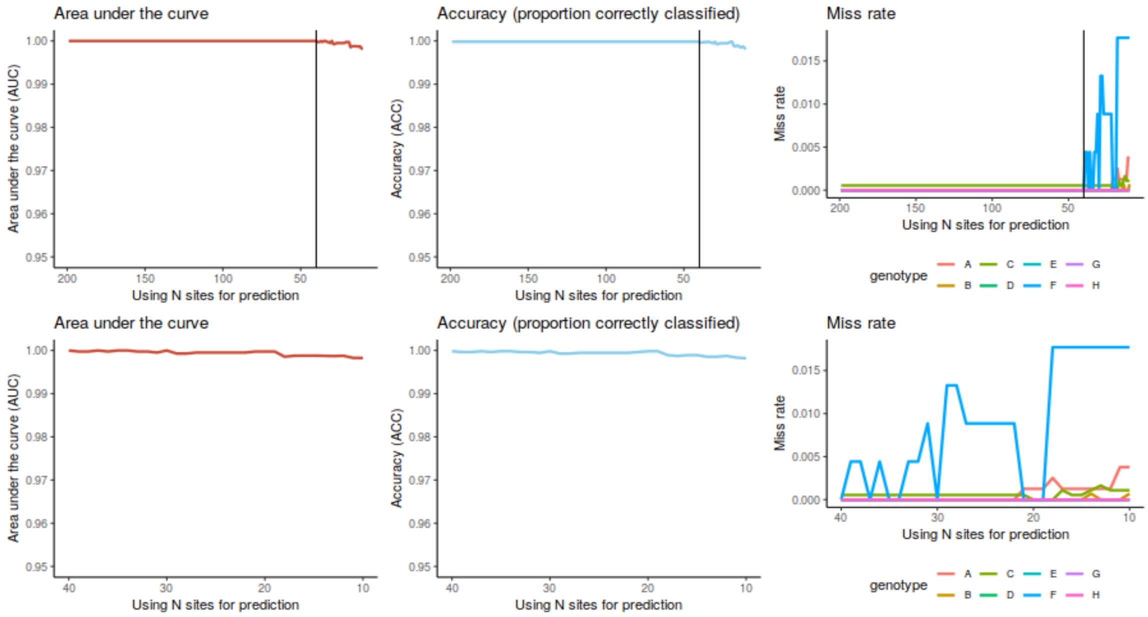

**Figure S5:** Phylogenetic trees showing (clockwise) the distribution of residues at sites 40 (HBx 40), 570 (Pol 201; spacer), 599 (Pol 230; spacer) and 637 (pol 268; spacer), all mapped by colour. Phylogenetic trees were available to download as a part of the online resource from which we obtained nucleotide sequences(4). Tree nodes where coloured on the basis of the amino acid residue at each site using an inhouse R script based on the R package Analyses of Phylogenetics and Evolution (ape v5.4 (12)). Only the top 5 most frequent amino acids are listed in the color legend, the rest are categorized as “other”. Phylogenies for the other 6 top-10 sites are shown in Figures 2 and S6.

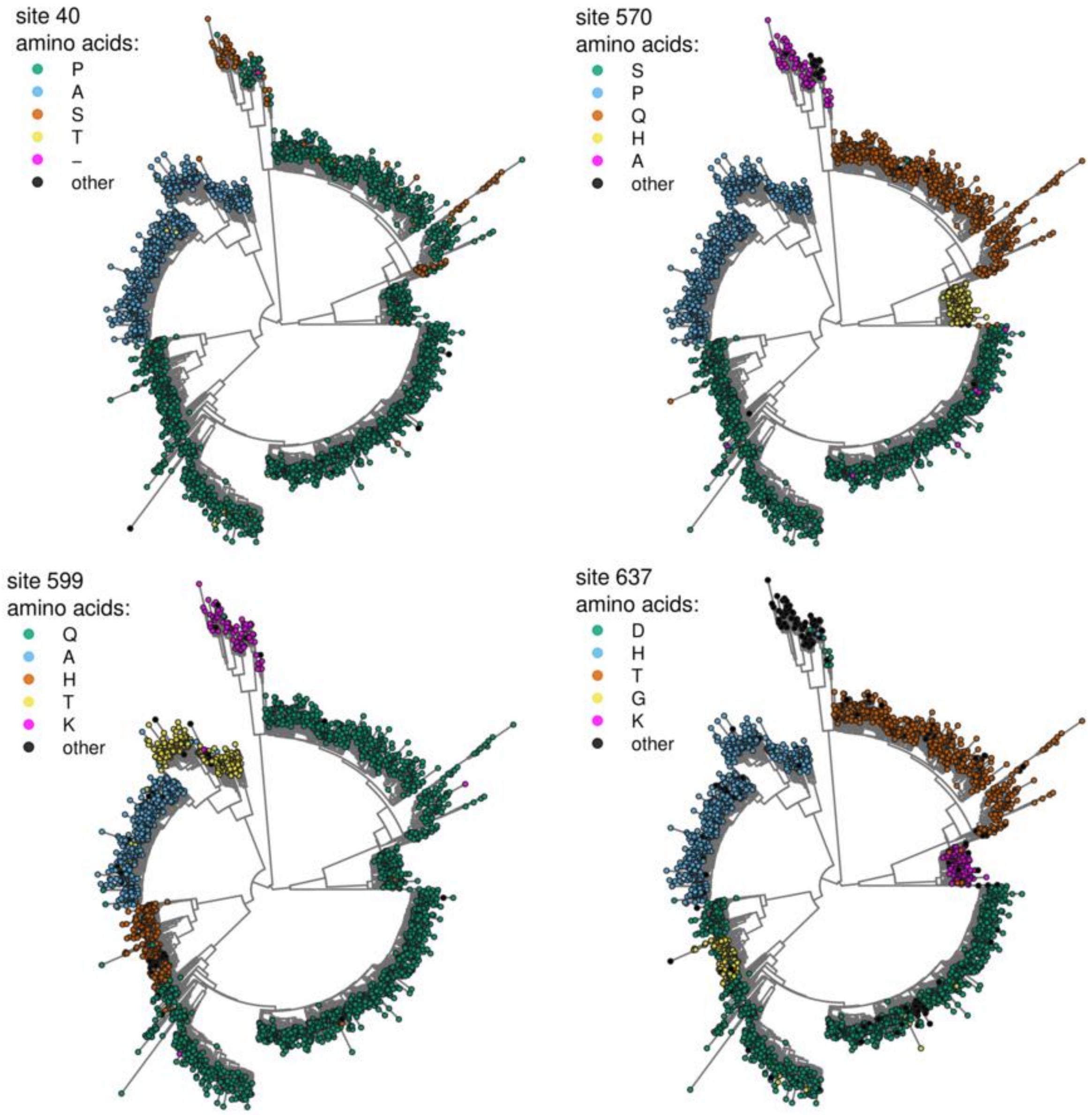

**Figure S6:** Phylogenetic trees showing (cloclwise) the distribution of residues at site 1253 (pre-S1 39),1262 (pre-S1 48) and 1435 (S 221), all mapped by colour. Phylogenetic trees were available to download as a part of the online resource from which we obtained nucleotide sequences(4). Tree nodes where coloured on the basis of the amino acid residue at each site using an inhouse R script based on the R package ‘Analyses of Phylogenetics and Evolution’ (ape v5.4 (12)). Only the top 5 most frequent amino acids are listed in the color legend, the rest are categorized as “other”. Phylogenies for the other 7 top-10 sites are shown in Figures 2 and S5.

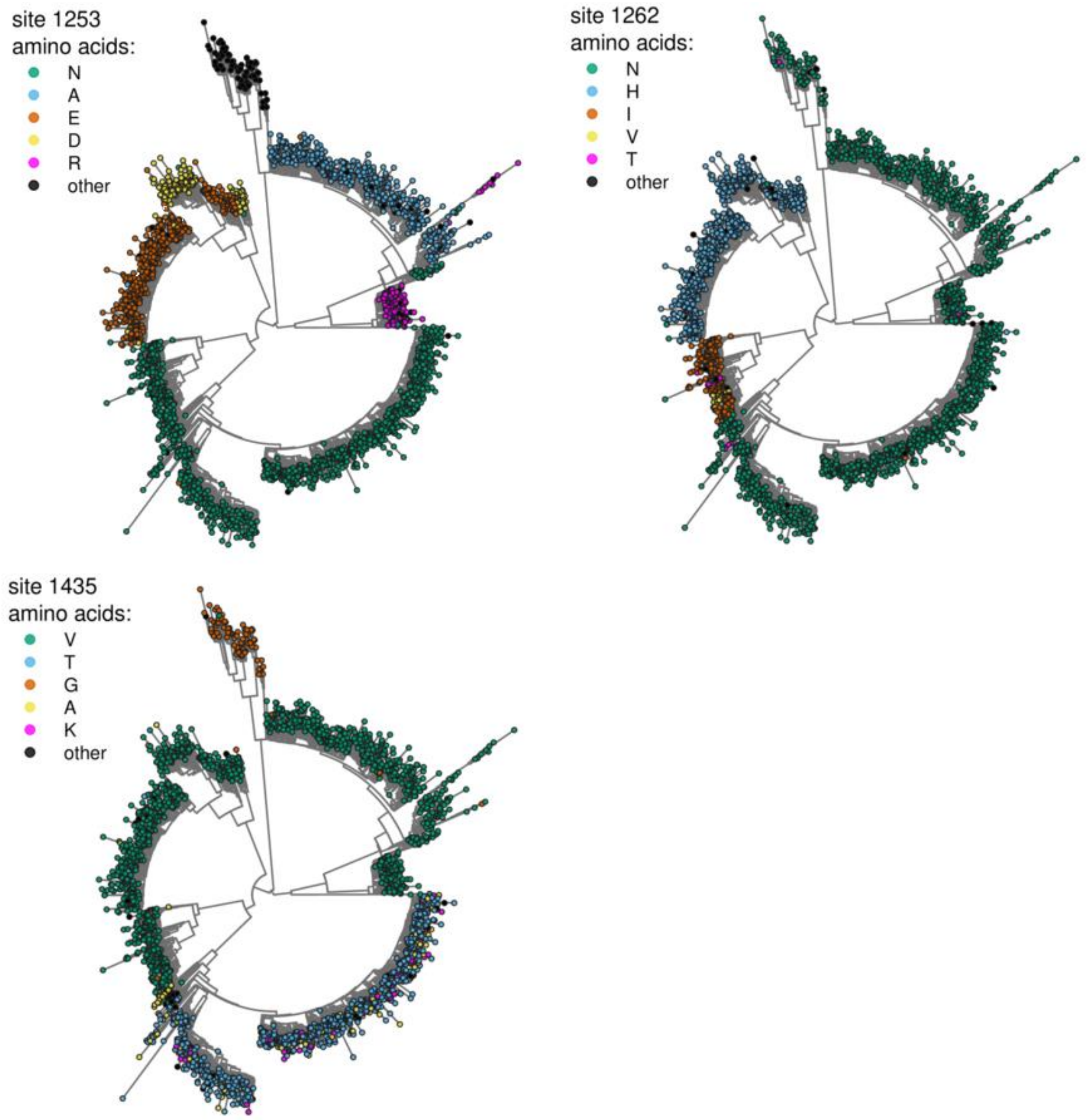

**Figure S7:** (A) Distribution of Mutual Information (MI) among all possible pairs of sites. The red vertical line marks MI=0.6, used as heuristic to evaluate pairs of sites with very high co-variation (MI>0.6). (B-D) Network of sites that show high co-variation with three sites in the top 10 list of sites most informative to predict genotype (center of networks). Link colours represent the level of MI between sites, with presented range of 0.6 – 0.89. The sites presented in panels (B-D) are listed in Table S4 with their respective MI values.

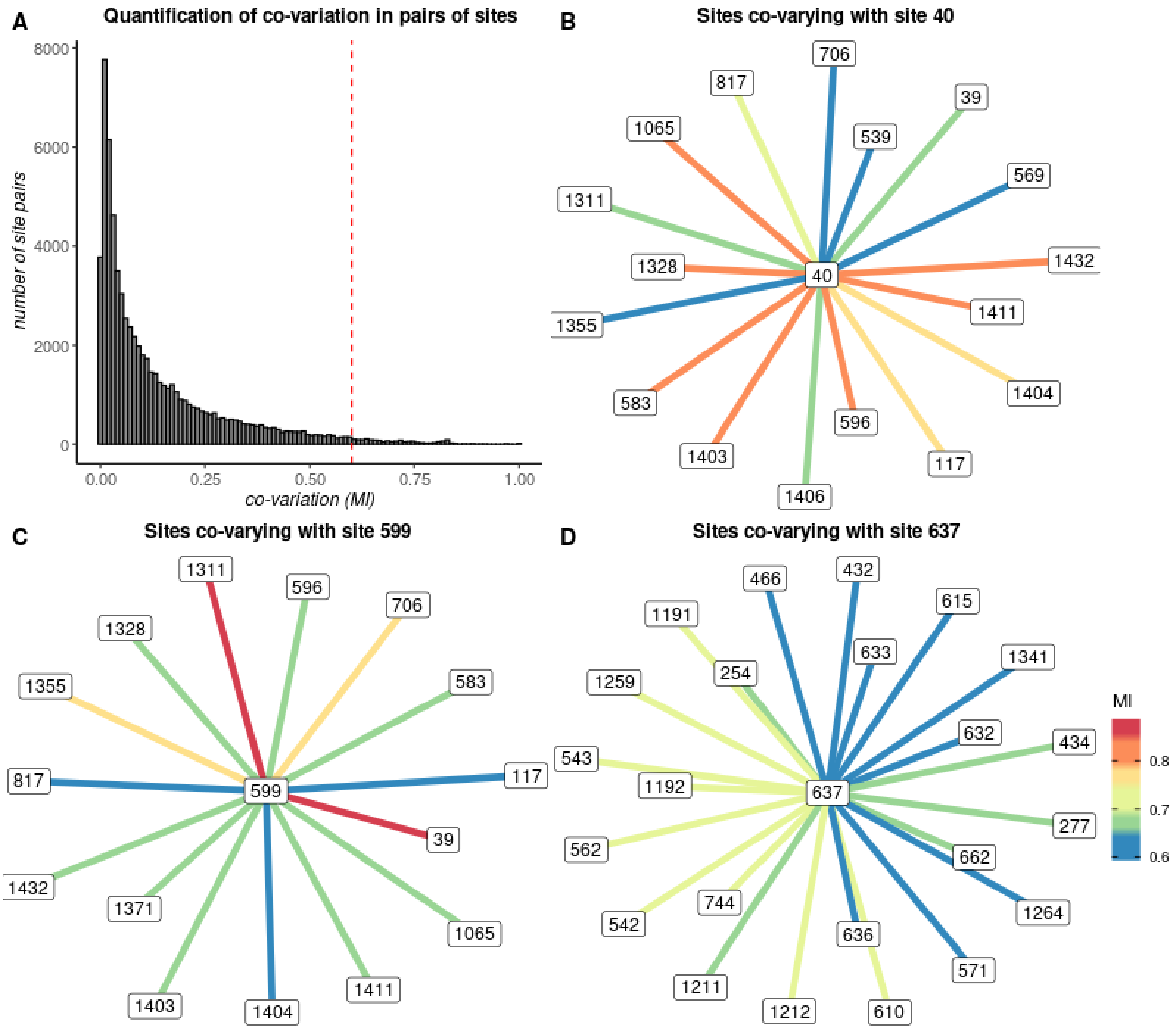

**Figure S8:** (A) Out-of-bag error distributions for predicting genotypes with sampling the dataset using a fixed proportion of samples per genotype, or (B) when sampling the dataset using a fixed number of samples per genotype. In both panels (A-B), results are presented when prediction is based on all amino acid sites (black) or the top 10 RFA selected sites (grey).

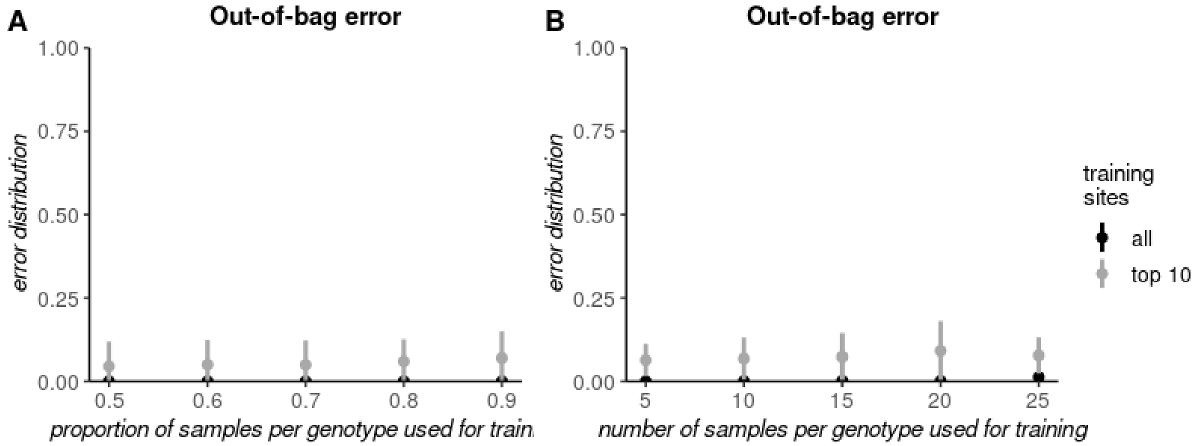
